# Beyond RNA Structure Alone: Complex-Aware Feature Fusion for Tertiary Structure-based RNA Design

**DOI:** 10.1101/2025.02.05.636748

**Authors:** Zixun Zhang, Jiayou Zheng, Yuzhe Zhou, Sheng Wang, Shuguang Cui, Zhen Li

## Abstract

Tertiary structure-based RNA design plays a crucial role in synthetic biology and therapeutics. While existing methods have explored structure-to-sequence mappings, they focus solely on RNA structures and overlook the role of complex-level information, which is crucial for effective RNA design. To address this limitation, we propose *t*the **C**omplex-**A**ware tertiary structure-based **R**NA **D**esign model, **CARD**, that integrates complex-level information to enhance tertiary structure-based RNA sequence design. To be specific, our method incorporates protein features extracted by protein language model (e.g., ESM-2), enabling the design model to generate more accurate and complex relevant sequences. Considering the biological complexity of protein-RNA interactions, we introduce a distance-aware filtering for local features from protein representation. Furthermore, we design a high-affinity design framework that combines our CARD with an affinity evaluation model. In this framework, candidate RNA sequences are generated and rigorously screened based on affinity and structural alignment to produce high-affinity RNA sequences. Extensive experiments demonstrate the effectiveness of our method with an improvement of **5.6%** compared with base model without our complex-aware feature integration. A concrete case study for 2LBS further validates the superiority of our CARD.

## 1. Introduction

RNA plays a pivotal role in biological research, serving as a key molecule in various cellular processes. Beyond its natural functions, RNA molecules design based on structures, also known as inverse RNA folding problem, has become increasingly important across diverse fields, including synthetic biology, therapeutics, and biotechnology. By leveraging the 3D structure, researchers can engineer RNA molecules to perform specialized and targeted functions, driving innovation in these areas.

RNA molecules design based on structure has been a long-standing focus in computational biology, driven by its applications in synthetic biology, therapeutics, and biotechnology. Early efforts rely on stochastic optimization and energy based optimization (Lorenz et al., 2011; Zuker, 2003) or physically informed optimization (Yesselman & Das, 2015). With the advancement of deep learning, this area has seen a shift toward more sophisticated computational approaches that can model the complex relationship between RNA sequences and their structures. For instance, LEARNA and Meta-LEARNA (Runge et al., 2018) leverages reinforcement learning for secondary structure-based design. While RDesign (Tan et al., 2024) and RhoDesign (Wong et al., 2024) utilize graph modeling to encode the 3D structures and decode the sequence using GNN and Transformer, respectively. RDesign (Tan et al., 2024) further leverages a contrastive learning to achieve data-efficient representation learning. RiboDiffusion (Huang et al., 2024) introduces the generative diffusion model for RNA inverse folding. However, these methods focus on modeling RNA structures alone, overlooking the crucial interaction dynamics in RNA complexes, as shown in Fig. 1 (a). As pointed out by Zhao (2024), different from protein folding which roughly follows Anfinsen’s rule, RNAs are highly flexible and rely on interactions with proteins, DNA, and other biomolecules to achieve folding. Furthermore, RNAs may adopt various conformations by interacting with distinct macromolecules at different stages of functioning. Given that RNA structures are heavily influenced by interactions within complexes, analyzing RNA structures alone is insufficient for inverse folding. A comprehensive approach must consider the complex-specific interactions that model RNA design and functionality.

**Figure 1.**
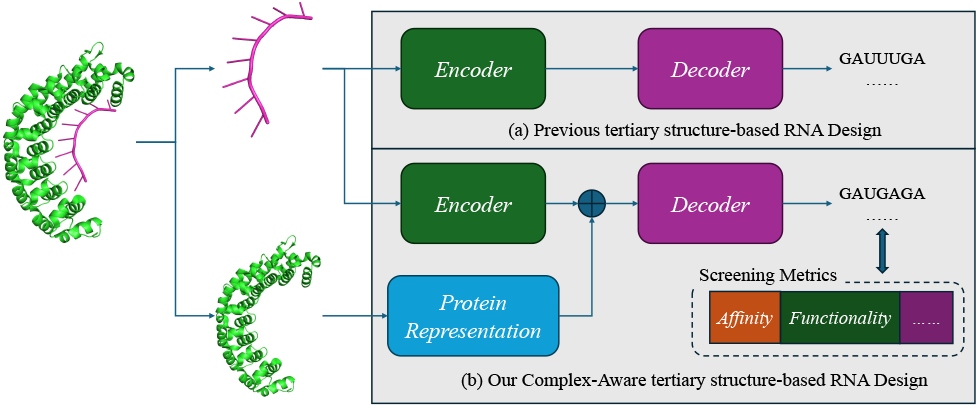
(a) Previous tertiary structure-based RNA design methods typically focus solely on RNA structural information. However, the critical interaction of RNA in complex has often been overlooked. (b) In this paper, we incorporate complex information into model to enhance its performance. Moreover, our model enables more flexible designs tailored to downstream requirements, such as focusing on functionality or affinity, thereby facilitating more targeted RNA design.

To address the above challenges, in this paper, we propose the **C**omplex-**A**ware tertiary structure-based **R**NA **D**esign model, **CARD**, that integrates RNA structural information and protein-RNA context to generate RNA sequences with complex context. In our CARD, apart from RNA tertiary structural representation, we utilize a pre-trained protein language model (PLM) to effectively represent the protein features, and fuse with the RNA representation to capture the complex interactions between RNA and proteins, as shown in Fig. 1 (b). Specifically, we adopt a Complex-Aware Transformer (CAFormer) to integrate RNA structural features and protein representations, enabling the generation of RNA sequences that adhere to key structural and complex context. Considering the biological reality that an RNA may interact with a protein through multiple binding sites and a single RNA can interact with different proteins, we introduce a distance-aware filtering to select the local representation for integration rather than using the representation of whole protein sequence. This filtering can make our model focus more on the interaction regions and discard the affect of irrelevant regions. By combining our design model with a learning based affinity prediction model, our framework can screen the generated RNA sequences through an iterative fashion. Candidate sequences are first filtered based on the predicted affinity scores to select the most promising candidates. These selected candidates are then validated for structural compatibility using multiple folding models, ensuring alignment with key structural metrics. By iteratively refining the candidate pool, our framework can ensure the designed sequences with both high affinity and structural compatibility, offering a computationally efficient alternative to traditional experimental methods like SELEX.

In summary, our contributions are three-fold:

1. We propose the Complex-Aware tertiary structure-based RNA design framework, **CARD**, that integrates RNA tertiary structures and protein-binding contexts, enabling a complex-aware RNA design.
2. For complex information fusion, we introduce the Complex-Aware Transformer (CAFormer) for complex information integration and a distance-aware filtering for the local features of protein representation, enabling the model to capture the protein-RNA complex context and focus on interaction regions within complex.
3. Extensive experiments on PRI30K and PRA201 (Han et al., 2024) datasets demonstrate the effectiveness of our framework. Additionally, a case study on high-affinity RNA design for 2LBS showcases the practical capacity and adaptability of our approach.

## 2. Related Work

### 2.1. Structure-to-Sequence RNA Design

RNA design aims to generate sequences that fold into pre-defined structures. Early methods focused on secondary structure optimization, using thermodynamic parameters and energy minimization. Tools like RNAfold (Lorenz et al., 2011) and Mfold (Zuker, 2003) predict RNA secondary structures based on the minimum free energy principle. As understanding of RNA structure has advanced, focus has shifted to complex tertiary structure-based design due to RNA’s high conformational flexibility, which challenges traditional thermodynamic methods (Ken et al., 2023). Recent deep learning approaches for RNA tertiary structure design include gRNAde (Joshi et al., 2024), which utilizes geometric deep learning and graph neural networks to generate RNA sequences; RiboDiffusion (Huang et al., 2024), a diffusion model for inverse folding leveraging RNA backbone structures; RDesign (Tan et al., 2024), which employs a data-efficient learning framework with contrastive learning for tertiary structures; and Rhodesign (Wong et al., 2024), focusing on RNA aptamer design by guiding sequence generation through structural predictions.

However, these methods often neglect the role of proteins in RNA design. Since RNA functionality depends not only on its structure but also on its interactions with proteins, we propose an RNA design approach that integrates protein information using protein language models.

### 2.2. Protein-RNA Complex Analysis

Protein-RNA interactions are crucial for gene regulation, RNA splicing, and stability. Various experimental techniques have been developed to characterize these interactions. Systematic Evolution of Ligands by EXponential Enrichment (SELEX) (Tuerk & Gold, 1990) is an in vitro method that selects high-affinity RNA aptamers for specific proteins. While useful for identifying RNA recognition motifs, it does not capture endogenous interactions in vivo. To address this, Crosslinking and Immunoprecipitation (CLIP) (Ule et al., 2005) employs UV crosslinking to stabilize RNA-protein complexes, followed by immunoprecipitation and sequencing. Variants such as HITS-CLIP (Licatalosi et al., 2008) and iCLIP (Konig et al., 2011) enhance resolution, while eCLIP (Van Nostrand et al., 2016) improves reproducibility and efficiency. RNA Immunoprecipitation Sequencing (RIP-seq) (Keene et al., 2006) provides a crosslink-free alternative, capturing RNA-protein interactions under physiological conditions, though with lower resolution. RNA Bind-N-Seq (RBNS) (Lambert et al., 2015) characterizes RNA-binding protein sequence preferences in vitro, revealing RNA recognition motifs.

However, these methods cannot be applied to proteins where experimental data does not exist. This limitation highlights the need for computational approaches capable of generalizing across different Protein-binding RNA.

### 2.3. Affinity Evaluation

The binding affinity between RNA and proteins is essential for validating RNA functionality. Current sequence-based prediction methods include PNAB (Yang & Deng, 2019), which manually extracts biochemical features and employs machine learning techniques like Support Vector Regression and Random Forests. DeePNAP (Pandey et al., 2024) utilizes convolutional neural networks to extract one-dimensional sequence features. Advanced approaches such as PredPRBA (Deng et al., 2019), PRdeltaGPred (Hong et al., 2023), and PRA-Pred (Harini et al., 2024) incorporate protein structural features—including secondary structures and binding interface characteristics like binding sites and hydrogen bonds—to achieve higher prediction accuracy with limited datasets.

However, the scarcity of RNA-protein complex structures presents significant challenges. To address this, CoPRA (Han et al., 2024) consolidates a protein-RNA binding affinity dataset with about 300 complexes, PRA310, from multiple sources and introduces a multi-modal model that integrates RNA and protein large language models with comprehensive structural information.

## 3. Method

The overall pipeline of our framework is shown in Fig. 2. Our complex-aware RNA design model, CARD, leverages the pre-trained protein language model to extract the representation of protein in the complex, and leverages CAFormer to integrate the complex information into RNA representation for more effective inverse folding. Accomplished with a machine learning based affinity prediction model and structural prediction tools, a pure computational RNA sequence screening framework can be obtained for both task-specific requirements and structural compatibility based on protein-RNA complexes. This framework provides a scalable and efficient alternative to experimental techniques like SELEX, enabling diverse applications such as therapeutic RNA design and functional genomics.

**Figure 2.**
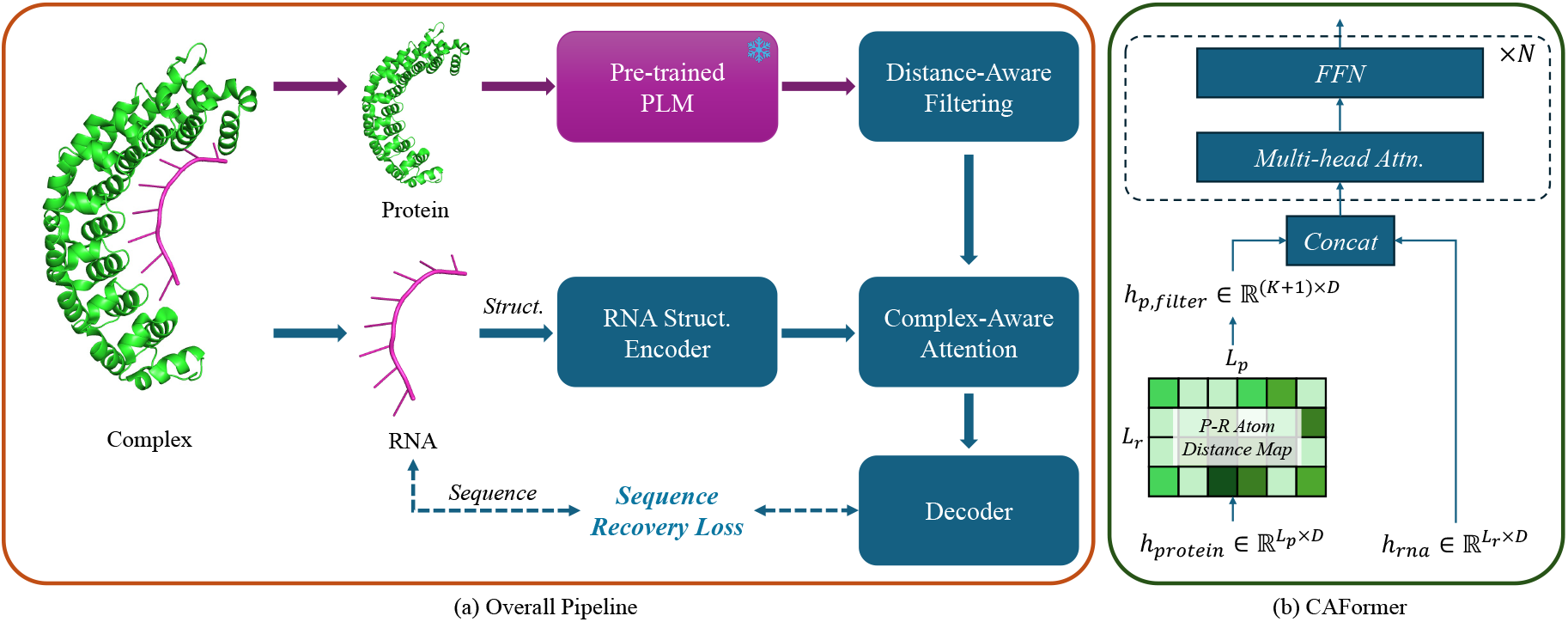
(a) The overall pipeline of our complex-aware tertiary structure-based RNA design model, CARD. We extract protein features by pre-trained PLM and integrate with RNA representation through CAFormer. (b) The detailed architecture of the CAFormer. The protein representation is first filtered via a distance-aware filtering to focus more on the protein-RNA interaction regions and then fused with the RNA feature through attention blocks.

### 3.1. RNA and Complex Representation Encoding

Given the protein-RNA complex, we first encode the RNA tertiary structures and protein representation, respectively. For the RNA structural representation, we utilize an architecture similar to RhoDesign (Wong et al., 2024), Geometric Vector Perceptron (GVP) (Jing et al., 2020), to encode the tertiary structures. To be specific, the GVP processes the RNA backbone coordinates (e.g., C4’, C1’, N1/N9) into vector and scalar features, integrating directional vectors and dihedral angles to capture structural information. This resulting representation, 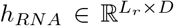 is utilized for complex-aware feature fusion with the protein features.

While for protein representation, protein sequences and structures are represented using pre-trained protein language models (PLMs), such as ESM-2 (Lin et al., 2022) or ESM-3 (Hayes et al., 2025). These PLMs can extract high-dimensional embeddings, 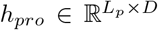, that capture sequence-level features, including secondary structure, functional motifs, and evolutionary relationships, providing a robust representation of the protein’s sequence and structural information.

### 3.2. Complex-Aware Feature Integration

After obtaining RNA structural representation, *h*_*RNA*_, and the representation of protein in complex, *h*_*pro*_, we incorporate Complex-Aware Transformer (CAFormer) blocks to integrate complex-specific information into the RNA representation, enabling more effective design. The CAFormer architecture consists of a distance-filtering block, followed by *N* Complex-Aware Attention blocks.

#### 3.2.1. Distance-aware Filtering

To make the feature fusion pay more attention to the information within binding regions of protein-RNA complex, we introduce a distance-filtering block. This block screens the protein representation, *h*_*P ro*_, based on the distances between protein amino acids and RNA nucleotides in the complex structure. Specifically, it selects *K* tokens corresponding to the closest amino acid-nucleotide pairs, as illustrated in Fig. 3. To preserve global information of the protein representation, we also concatenate a global protein representation token by global average pooling (GAP) with the locally selected tokens, resulting in the filtered protein representation, *h*_*pro,filtered*_ *∈* ℝ^*K*+1*×D*^.

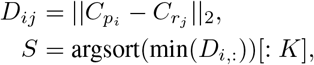

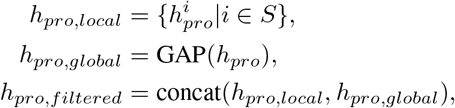

where 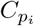 and 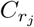 are the coordinates of the center of the *i*-th amino acid and *j*-th RNA nucleotide, respectively.

**Figure 3.**
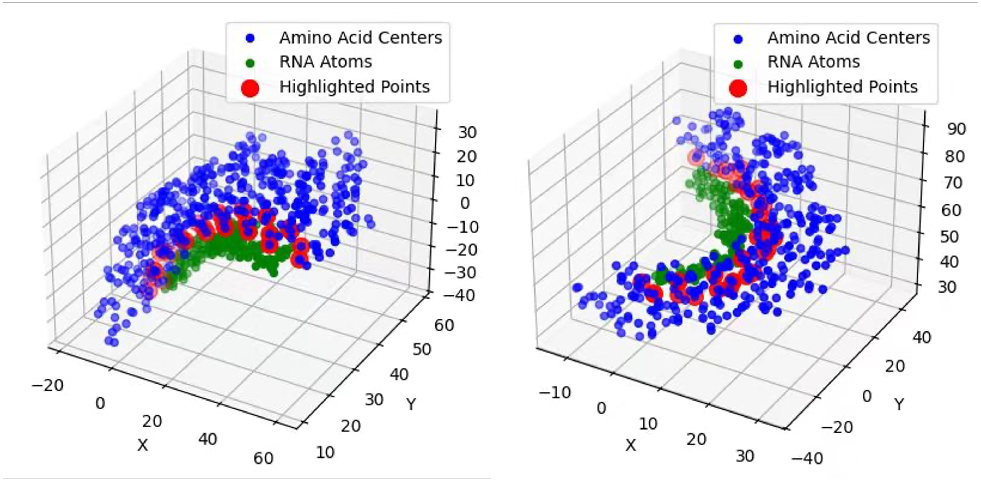
An example of distance-aware filtering. Based on the distance between amino acids and nucleotides, we select the nearest *K* amino acids to construct the protein representation, the selected indices are highlighted in red in the figure.

Notably, considering the biological complexity of protein-RNA interactions, where a protein may bind to an RNA through multiple binding sites (different poses), and also a single RNA can interact with different proteins, this distance-aware filtering not only captures the global information of the protein within the complex but also enables the model to focus more on the features of the interaction regions. Fig. 4 gives two examples of protein-RNA multi interaction of a single RNA with the same protein and different proteins. By leveraging this filtering, we can integrate complex features more efficiently while mitigating the influence of non-interaction regions, thereby obtaining a more refined RNA representation.

**Figure 4.**
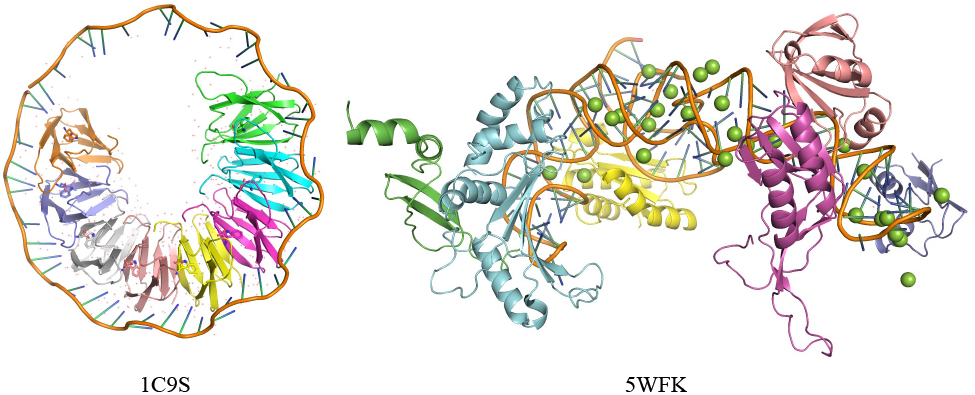
An example of protein-RNA multi interaction from 1C9S and 5WFK. Better viewing with color mode.

#### 3.2.2. Complex-AwareAttentionBlock

Given the RNA structural representation, *h*_*RNA*_ and the filtered protein representation, *h*_*pro,filtered*_, we first concatenate these two representations to form a unified complex representation, 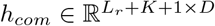. To incorporate complex-specific information and refine the RNA representation, we leverage *N* multi-head self-attentions with *h*_*com*_ as key and value, enabling the RNA representation to capture the relevant features from the protein context. This process enhances the RNA representation with complex-aware information, facilitating more effective RNA design.

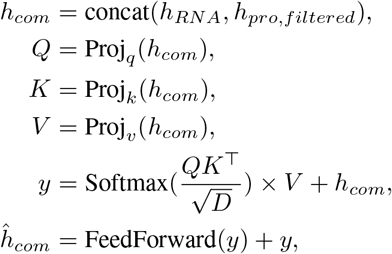

where Proj_*_ represents the linear projection for *Q, K*, and *V*. For simplicity, here we omit the LayerNorm operations and output projection of multi-head attention. After applying the complex-aware attention, we extract the indices corresponding to the RNA representation, which serve as the final RNA representation for decoding operations.

For decoding enhanced RNA representation, we employ a transformer-based decoder similar to RhoDesign (Wong et al., 2024) to generate RNA sequences. The decoding process is formulated as a next-token prediction task, where the decoder predicts the next nucleotide based on preceding tokens. During training, the model is trained with teacher forcing using CE loss.

### 3.3. High-Affinity RNA Design Framework

Building upon the inverse folding model detailed above, in this section, we present the high-affinity RNA Design framework, a comprehensive framework that integrates protein-RNA complex information and evaluation mechanisms. This framework is to design RNA sequences optimized under the constraint of high binding affinity and structural compatibility, ensuring suitability for diverse downstream applications. The design framework consists of two main components: the structure-to-sequence design model and evaluation tools.

#### Design Phase

In this stage, our complex-aware design model synthesizes candidate RNA sequences by integrating RNA tertiary structures with protein-RNA complex information. Utilizing pre-trained protein language models (PLMs) and our CAFormer, the design model can effectively capture the intricate relationships between RNA and binding proteins, enabling the generation of RNA sequences tailored for complex constraints, such as binding sites and interactions.

#### Constraint Evaluation

Following the sequence generation, the constraint evaluation is utilized to filter the RNA sequences to meet the task-specific requirements while maintaining structural compatibility within the protein-RNA complex. This evaluation combines metrics for both affinity evaluation and structural similarity. For the affinity evaluation, we train an ensemble regression model on the PRA201 (Han et al., 2024) dataset to predict the affinity score based on protein and RNA sequences. While for structural evaluation, we calculate RMSD score and TM-Score by aligning the predicted structures with the original ones. To mitigate the potential errors from low-confidence structural predictions, we employ multiple structure prediction models, including AlphaFold3 (Abramson et al., 2024), RhoFold (Shen et al., 2024), and RoseTTAFold2NA (Baek et al., 2024), etc. It is worth noting that the affinity evaluation can be replaced with other functional metrics to adapt to different task-specific requirements, providing flexibility for various applications.

#### Iterative Screening

To achieve high-throughput screening while balancing computational efficiency, the searching process adopts a two-step evaluation. In each iteration, the pipeline first evaluates the binding affinity of the RNA candidates, selecting the top 10% ∼ 20% of sequences based on their affinity scores. These selected candidates are then sent to folding models for structural validation to obtain the most optimal candidates. The optimal candidates obtained will serve as templates for the following iterations, where new RNA sequences are generated and re-evaluated through the same process. By iteratively updating the candidate pool, the framework ensures that the generated RNA candidates not only exhibit high affinity but also maintain structural compatibility by pure computational tools, which can offer the potential to assist or even replace time- and labor-intensive experimental techniques like SELEX in the field of RNA design.

## 4. Experiment

### 4.1. Dataset and Implementation Details

#### Dataset

To evaluate the effectiveness of our CARD, we curate our dataset from the PRI30K and PRA201 datasets proposed in Han et al. (2024). PRI30K is a non-redundant collection of protein-RNA interaction pairs, encompassing proteins with a maximum residue length of 750 and RNA sequences ranging from 5 to 500 bases (5 ≤ *L*_*r*_ ≤ 500). While PRA201 is compiled from PDBbind (Wang et al., 2004), PRBABv2 (Hong et al., 2023), and ProNAB (Harini et al., 2022), encompassing 201 unique protein-RNA complexes. Each sample comprises a single protein chain and a single RNA chain.

Here we directly utilize PRA201 as the blind test set for evaluation and downstream high-affinity RNA design studies. While for PRI30K, we first exclude the RNA chains having high similarity with those in PRA201 and over-length RNA chains for computational efficiency (*L*_*r*_ *>* 128), resulting in a dataset comprising 21,050 protein-RNA pairs and 2,309 unique RNA sequences. For unbiased model evaluation, we utilize **CD-HIT** (Li & Godzik, 2006) for clustering all RNA chains based on nucleotide sequence similarity. Specifically, RNA sequences exhibiting ≥80% sequence identity are grouped into the same cluster. Following this clustering, the dataset is randomly divided into training and test sets in an 80%-20% ratio.

#### Training Details

We train our model on two A100 GPUs for 200 epochs with a batch size of 48 (24 per GPU) using our training split of PRI30K. In CAFormer, we set the number of attention blocks to *N* = 6 and the local filtered size of the protein representation to *K* = 64. The protein representation is extracted by ESM-2 650M. The total training time is approximately 6 hours.

### 4.2. Quantitative Comparison of Complex-Aware RNA Design

We evaluate our method on the two test sets and compare it with several models, including two sequence-based baselines (SeqRNN and SeqLSTM), StructGNN, GraphTrans (Ingraham et al., 2019), RDesign (Tan et al., 2024), and RhoDesign (Wong et al., 2024).

#### PRI30K Test Split

Tab. 1 presents the comparison results on the split test set of PRI30K. Our method achieves consistent and significant improvements over all reproduced baselines. Our method achieves the highest recovery rate and macro F1 score among all length categories with an overall recovery rate of 59.56% and a macro F1 score of 0.6002. In particular, traditional sequence-based methods perform poorly, where SeqLSTM with a hidden size of 128 achieves a recovery rate of only 29.00%, 31.28%, and 30.94% on short, medium, and long sequence categories, which are slightly better than random guess. Increasing the hidden size provides marginal improvement, but overall, both SeqLSTM and SeqRNN are outperformed by our method by about 30% in all sequence categories. While for StructGNN and GraphTrans which are designed for protein inverse folding, they achieve an overall recovery rate of about 37%. Compared to RhoDesign, which can be seen as a base model of our method, we provide an improvement of 2.78%, 5.2%, and 10.32% on short, medium, and long sequence categories, respectively. Our method exhibits a similar trend in Macro F1 performance. Compared to the second-best method, RhoDesign (Wong et al., 2024), our method achieves consistent improvements across all sequence length categories, with gains ranging from 0.04 to 0.1. Specifically, our model improves the overall macro F1 score by 0.06. The overall results reinforces the effectiveness of our method of capturing complex-aware feature into RNA design, leading to more reliable predictions.

**Table 1.** Comparison results on the test set of PRI30K. ^*\**^: We reproduce all the other methods on our PRI30K training set except for RDesign, due to the unavailability of their training code or insufficient algorithmic details. Therefore, we directly utilized the public checkpoint.

| Method | Recovery Rate (%) |  |  |  | Macro F1 |  |  |  |
| --- | --- | --- | --- | --- | --- | --- | --- | --- |
|  | Short | Medium | Long | All | Short | Medium | Long | All |
| SeqLSTM(h=128) | 29.00% | 31.28% | 30.94% | 30.65% | 0.2814 | 0.3166 | 0.3012 | 0.2927 |
| SeqLSTM(h=256) | 29.02% | 31.04% | 29.26% | 30.16% | 0.2867 | 0.3117 | 0.2870 | 0.2935 |
| SeqRNN(h=128) | 28.00% | 29.14% | 30.29% | 29.11% | 0.2731 | 0.2967 | 0.2996 | 0.2819 |
| SeqRNN(h=256) | 29.22% | 31.45% | 31.26% | 30.86% | 0.2892 | 0.3208 | 0.3006 | 0.2988 |
| GraphTrans | 31.22% | 39.69% | 41.15% | 37.17% | 0.3136 | 0.3945 | 0.4101 | 0.3430 |
| StructGNN | 30.89% | 39.53% | 42.19% | 37.22% | 0.3061 | 0.3933 | 0.4206 | 0.3386 |
| RhoDesign | 56.18% | 54.92% | 49.07% | 54.00% | 0.5618 | 0.5492 | 0.4907 | 0.5400 |
| RDesign* | 36.19% | 49.00% | 52.40% | 45.64% | 0.3574 | 0.4872 | 0.5224 | 0.4070 |
| CARD (Ours) | **58.96%** | **60.12%** | **59.39%** | **59.56%** | **0.6009** | **0.6007** | **0.5938** | **0.6002** |

#### PRA201 Blind Test

Tab. 2 presents a quantitative comparison of different methods on the PRA201. Our method consistently outperforms other methods on both recovery rate and macro F1 score. Compared to the second-best method, RhoDesign (Wong et al., 2024), we achieve an improvement of 6.21%, 11.19%, and 7.59% of recovery rate, respectively. Similarly, our method achieves the best overall macro F1 score of 0.6108, outperforming the second-best RhoDesign by 0.0431. This result further enhances the effectiveness of the design of our complex-aware attention with distance-aware filtering mechanisms.

**Table 2.** Comparison results on the PRA201.

| Method | Recovery Rate (%) |  |  |
| --- | --- | --- | --- |
|  | Short | Medium | All |
| GraphTrans | 30.73% | 43.47% | 33.33% |
| StructGNN | 30.94% | 41.16% | 33.33% |
| RhoDesign | 55.95% | 57.54% | 56.39% |
| RDesign* | 36.50% | 52.31% | 40.88% |
| CARD (Ours) | **62.16%** | **68.73%** | **63.98%** |

| Method | Macro F1 |  |  |
| --- | --- | --- | --- |
|  | Short | Medium | All |
| GraphTrans | 0.2940 | 0.4351 | 0.3020 |
| StructGNN | 0.3046 | 0.4131 | 0.3107 |
| RhoDesign | 0.5674 | 0.5722 | 0.5677 |
| RDesign* | 0.3537 | 0.5218 | 0.3658 |
| CARD (Ours) | **0.6051** | **0.6830** | **0.6108** |

Fig. 5 gives a violin plot of the detailed distribution of recovery rates for different methods across two test sets. It can be observed that GraphTrans (Ingraham et al., 2019), StructGNN, and RDesign (Tan et al., 2024) exhibit wider and more dispersed distributions with a lower median recovery rate, indicating high variance and inconsistent performance across different sequence lengths. RhoDesign (Wong et al., 2024) demonstrates a more stable performance with a slightly higher median recovery rate. Overall, our method maintains a clear advantage over other methods. While other methods exhibit wider spread and lower overall recovery rates, our method produces a more compact and higher-distribution shape, demonstrating better stability and reliability performance across different datasets.

**Figure 5.**
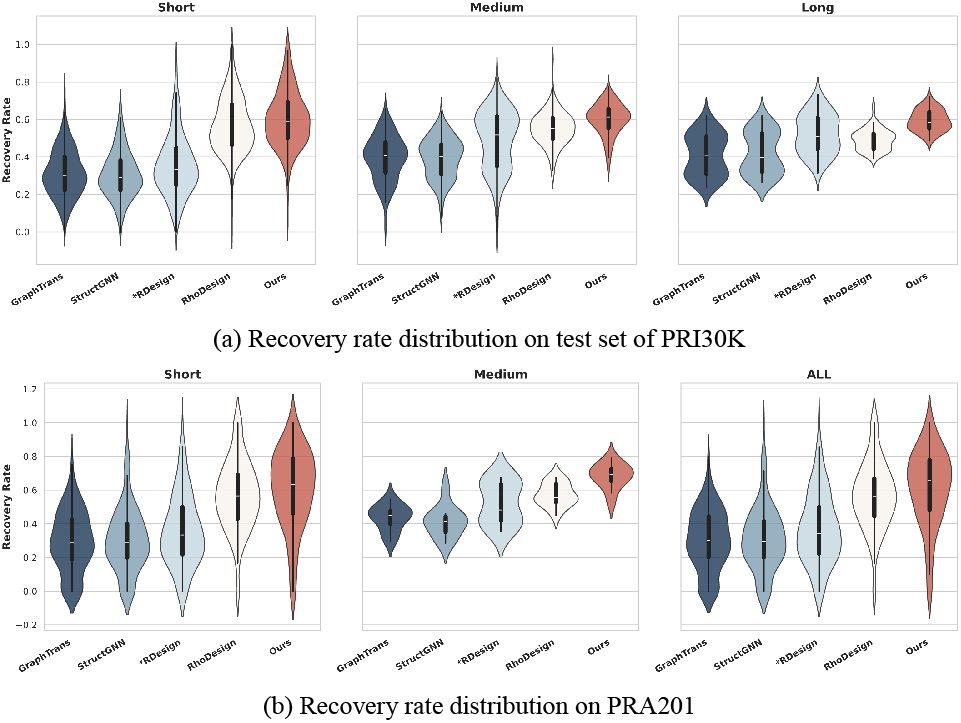
Violin plots for the recovery rate distribution of different methods across datasets and sequence categories.

### 4.3. Ablation Studies

In this section, we conduct several detailed analysis of the key components of our method, including the number of attention blocks in CAFormer, the fashion of filtering the protein representation, and the choices of protein representation model. All the models are evaluated on PRA201.

#### 4.3.1. Impact of Block Numbers of CAFormer

In this part, we conduct several experiments to figure out the impact of numbers of complex-aware attention blocks as shown in Tab. 3. Without any complex-aware attention blocks where the design model only consists of RNA structure encoder and decoder blocks, the overall macro F1 score is 0.5677. With two blocks of CAFormer, the score achieves an improvement of 0.0322, 0.0971, and 0.0369 in short, medium length categories, and overall macro F1 score. This improvement demonstrate the effectiveness of integrating complex information into RNA design. The performance consistently improves as the blocks increasing. Our method achieves the highest macro F1 score with six blocks across different length categories.

**Table 3.**
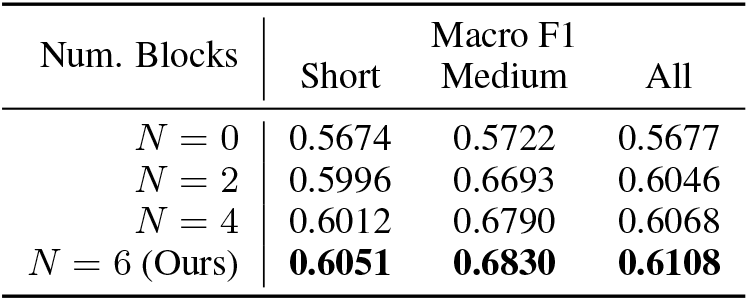
Ablation of number of blocks in CAFormer.

| Num. Blocks | Macro F1 |  |  |
| --- | --- | --- | --- |
|  | Short | Medium | All |
| $N = 0$ | 0.5674 | 0.5722 | 0.5677 |
| $N = 2$ | 0.5996 | 0.6693 | 0.6046 |
| $N = 4$ | 0.6012 | 0.6790 | 0.6068 |
| $N = 6$ (Ours) | **0.6051** | **0.6830** | **0.6108** |

#### 4.3.2. Impact of Protein Feature Filtering

To figure out the best fashion of protein feature integration, we conduct several experiments as shown in Tab. 4. With the global representation by global average pooling of protein only, the model achieves an improvement of 0.0232 on overall macro F1 score. This result further reinforce the effectiveness of integration protein-RNA interaction context into RNA design model. While applying random selection for local representation, the performance degrades significantly, decreased by 0.0193 and 0.0163 on short sequence category and overall score, respectively. With our distance-aware filtering, the performance further achieves an improvement of 0.0176, 0.0485, and 0.0199, respectively. The experimental results validate the effectiveness of our distance-based filtering for protein-RNA interaction regions. Our distance-aware approach efficiently filters out irrelevant regions while preserving the essential interaction context.

**Table 4.** Ablation of the fashion of complex feature integration. GR denotes global representation of protein. LR represents the local representation of protein. Rand. and Dis. are random selection and our distance-aware filtering, respectively.

| Global | Local | Macro F1 |  |  |
| --- | --- | --- | --- | --- |
|  |  | Short | Medium | All |
| ✓ | ✗ | 0.5875 | 0.6345 | 0.5909 |
| ✓ | Random | 0.5682 | 0.6568 | 0.5746 |
| ✓ | Distance | **0.6051** | **0.6830** | **0.6108** |

#### 4.3.3. Choices of Protein Representation

In this part, we further conduct experiments using different size of ESM-2 to figure out the impact of protein representation models. The results are shown in Fig. 6. The impact of the ESM-2 model size on performance is relatively minor. Using 650M model for protein representation achieves an average improvement of only 1.25% in recovery rate and an average increasement of 0.0041 in macro F1 score compared to the 35M model.

**Figure 6.**
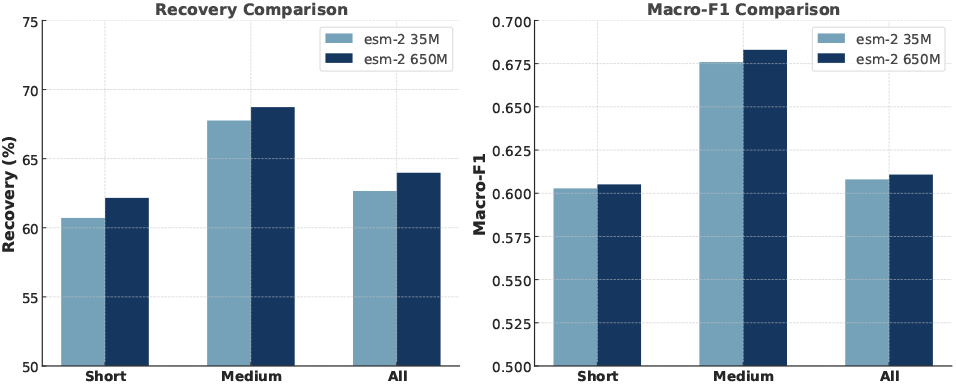
Performance of complex-aware attention integrating with representation produced by different ESM-2 models.

## 5. High-Affinity RNA Design for 2LBS

In this section, using 2LBS as an example, we demonstrate how our CARD can enhance the binding affinity of RNA to its target protein while maintaining the stability of the RNA structure using the high-affinity design framework described in Sec. 3.3. 2LBS is the solution structure of the double-stranded RNA binding domain of S. cerevisiae RNase III (Rnt1p) in complex with the AAGU tetraloop hairpin.

We first input the native 2LBS complex into our CARD and perform RNA inverse folding to generate 1000 different RNA sequences. We then predict the binding affinities of these RNA sequences and select the top 20 sequences with the highest affinity. Next, we use AlphaFold3 (Abramson et al., 2024) to calculate the complex structures of these high-affinity RNAs with the target protein, and use these structures as inputs for the second round of design. For each complex, we generate 50 RNA sequences, resulting in a total of 1000 RNA sequences for the second round.

This process is repeated to complete the third round of design. In Fig. 7, we present two RNA sequences from our design that exhibit higher binding affinity than the native RNA sequence, with structural differences from the native sequence being minimal. We present the distribution of the binding affinity of designed RNA sequences in Fig. 8. This further validates the effectiveness of the CARD method in optimizing RNA binding affinity. More details are provided in the appendix.

**Figure 7.**
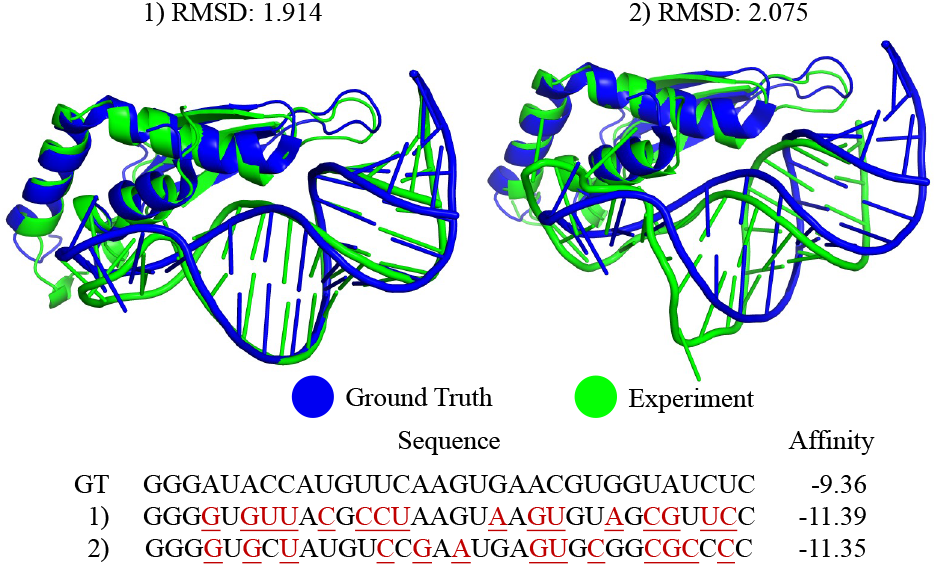
Structures and sequences of RNAs designed by CARD for 2LBS. The structures are predicted by AlphaFold3 and aligned with the naive structure.

**Figure 8.**
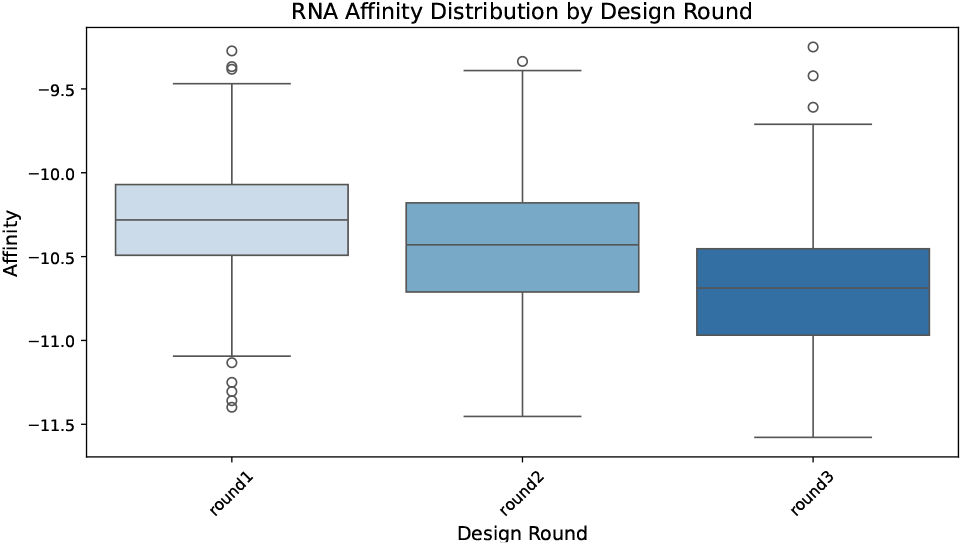
The distribution of predicted affinity across 3 design round.

## 6. Conclusion

In this paper, we propose a complex-aware tertiary structure-based RNA design framework, CARD, that integrates RNA structure representations and protein-binding contexts. Unlike existing approaches that focus solely on RNA structure, our framework explicitly incorporates protein-RNA interactions into the inverse folding process. To achieve this, we introduce the Complex-Aware Transformer (CAFormer) to fuse RNA and protein representations, ensuring RNA sequences adhere to both structural and complex constraints. Additionally, to make the model focus more on the interaction regions, we design a distance-aware filtering to select the local protein representations for fusing rather than the whole protein representations. Our method further incorporates an iterative screening process that integrates an affinity predictor and structural validation, enabling efficient and scalable optimization of RNA sequences. Extensive experiments on the PRI30K and PRA201 datasets demonstrate the effectiveness of our approach. By integrating RNA tertiary structure modeling with protein-RNA interactions, our framework provides a robust and computationally efficient solution for RNA sequence design exploring the sequence space under diverse constraints.

### Impact Statements

The goal of our paper is to advance the computational RNA structure-aware design via deep learning approaches. Computational design approach can advance the development of targeted RNA drugs, vaccines, and gene therapies, contributing to personalized medicine and precision therapeutics. This could also streamline synthetic biology workflows, enabling automated discovery of functional RNAs for biosensors, riboswitches, and programmable gene circuits. However, our method relies on large-scale biological datasets for training, which may contain biases due to limited experimental data availability. Ensuring diverse and high-quality datasets is crucial to prevent model biases that could impact prediction or design. Furthermore, our method is only evaluated on public datasets and has not been tested experimentally in real biological scenarios which is critical for real-world applications.

## Acknowledgements

This work was supported by Shenzhen-Hong Kong Joint Funding No. SGDX20211123112401002, by the Basic Research Project No. HZQB-KCZYZ-2021067 of Hetao Shenzhen HK S&T Cooperation Zone, by Shenzhen General Program No. JCYJ20220530143600001, by the Shenzhen Outstanding Talents Training Fund 202002, by Guangdong Research Project No. 2017ZT07×152 and No. 2019CX01×104, by the Guangdong Provincial Key Laboratory of Future Networks of Intelligence (Grant No. 2022B1212010001), by the Guangdong Provincial Key Laboratory of Big Data Computing, CHUK-Shenzhen, by the NSFC 61931024&12326610, by the Key Area R&D Program of Guangdong Province with grant No. 2018B030338001, by the Shenzhen Key Laboratory of Big Data and Artificial Intelligence (Grant No. ZDSYS201707251409055), and by Tencent & Huawei Open Fund.

## A. Dataset Details

As mentioned in the main paper, we filter out the protein-RNA pairs from PRI30K to train our CARD. Fig. 9 shows the detailed distribution of our filtered sequences. The left part shows the frequency distribution of RNA sequence lengths in the dataset. The average sequence length is 42.9 with a STD of 33.7. Two peak regions can be observed in the length distribution, a primary peak in 0 *∼* 30 nucleotide range and a secondary peak in the 60 *∼* 80 nucleotide range.

**Figure 9.**
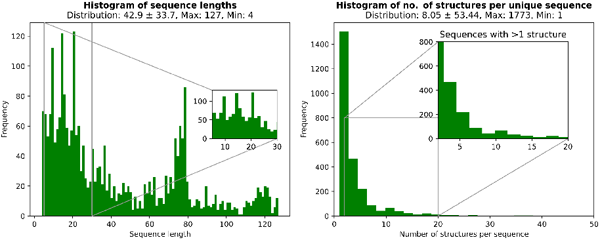
The detailed distribution over sequence length and protein-RNA pairs.

While the right part of the figure shows the histogram of protein-RNA pair counts per unique RNA sequence. This histogram exhibits a typical long-tail distribution, where the majority of RNA sequences are associated with only a small number of protein-RNA interaction structures, while a few sequences have an exceptionally large number of structures. The maximum number of structures per sequence is 1773.

## B. More Details of Case Study for 2LBS

In this part, we show more details of the case study for 2LBS. Fig. 10 shows a sequence logo of an RNA sequence generated using Logomaker, which visually represents the composition and positional conservation in the RNA sequence. The vertical axis shows the frequency of {A, U, C, G} at each position, while the height of each letter corresponds to its occurrence. It can be observed that certain positions are dominated by a relatively deterministic nucleotide, such as G at position 0 ∼ 3, GUCUGA at position 10 ∼ 15, and GUCCC at the end of the sequence. The conserved regions likely play essential roles in RNA stability or interactions, while the more variable positions suggest flexible or adaptive functional sites. The similar phenomenon is also reflected in Fig. 7 of the main text. The non-highlighted positions in the sequence and the interface regions in the folding visualization exhibit a relatively strong resemblance to the naive structure, indicating that these regions remain structurally stable.

**Figure 10.**
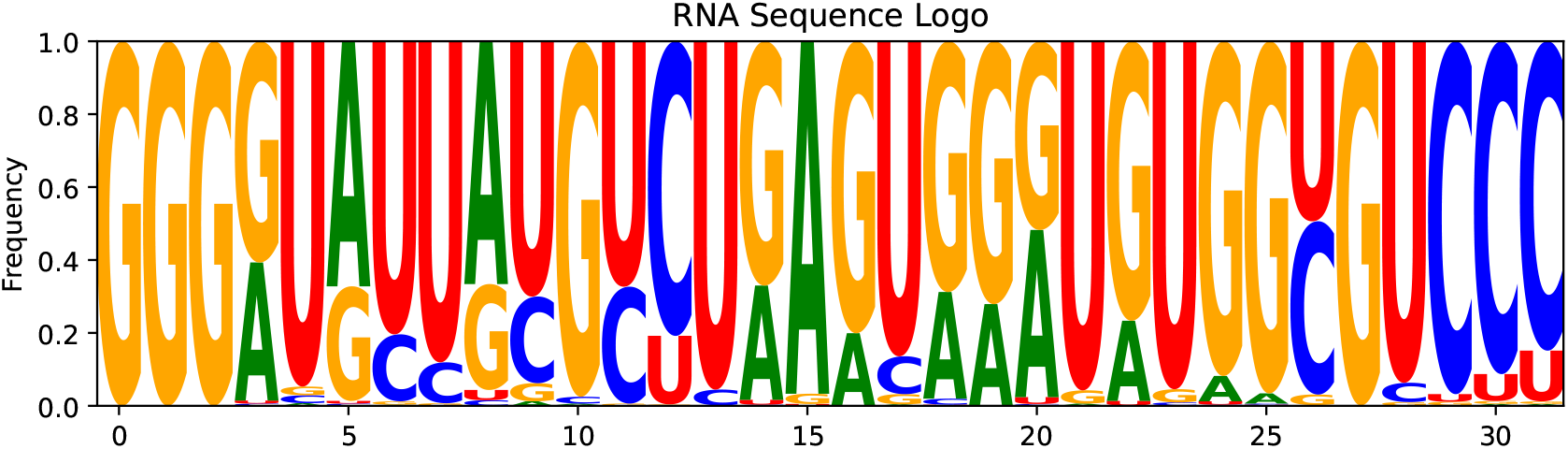
A sequence logo of generated RNA sequences.

In Fig. 11, we further show the heatmaps of predicted chemical modification reactivity under 2A3 and DMS. Both 2A3 reactivity and DMS reactivity indicate that strong high-reactivity regions (deep red) are clustered in the central segment, particularly within the loop regions. The designed sequence and groundtruth align well, showing that our CARD accurately captures chemical modification sensitivity.

**Figure 11.**
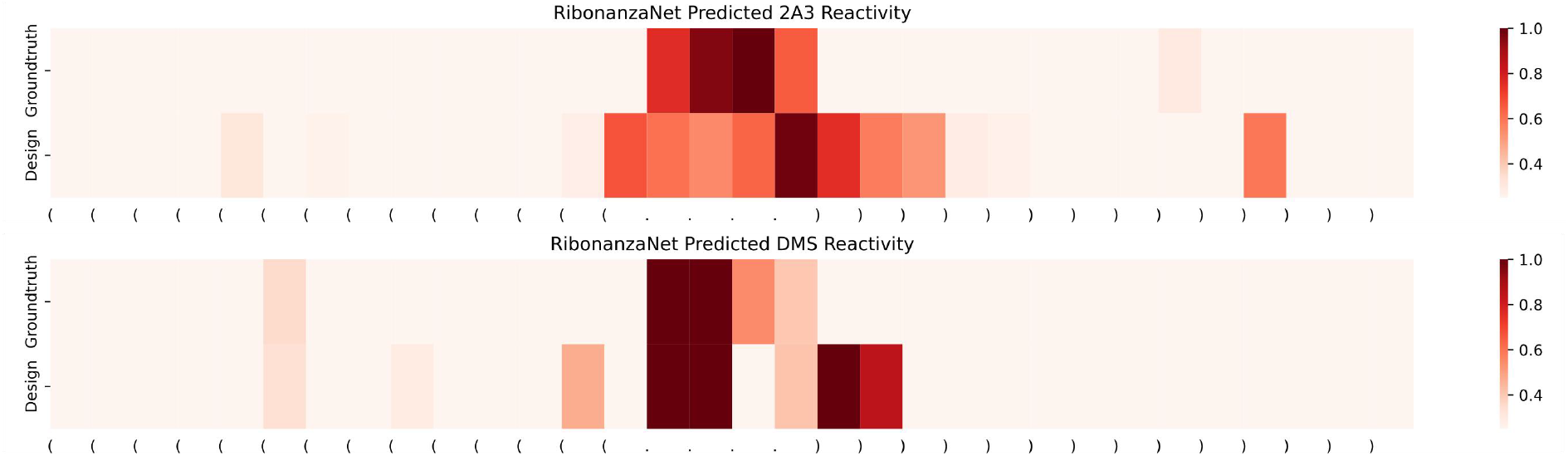
The detailed distribution over sequence length and protein-RNA pairs.

## C. Affinity Prediction Model

As mentioned in the main text, we leverage a learning based affinity prediction model for high-throughput screening. In this part, we present how we conduct our affinity prediction model. Although CoPRA (Han et al., 2024) introduces an affinity prediction model by combining protein and RNA language models, it relies heavily on structural information, which contradicts our goal of achieving high-throughput screening. Furthermore, the structure predicted by folding models is often inaccurate, which can further negatively impact affinity predictions that depend on structural features.

To this end, we design a concise pure sequence based affinity predictor. Similar to CoPRA, we first extract protein sequence representation and RNA sequence representation by ESM-2 and RiNaLMo, and obtain the global representation via global average pooling. We then concatenate them to form the overall sequence feature *h ∈* ℝ^2560^. While for the regression model, we employ an ensemble approach that integrates three models: LightGBM, XGBoost Random Forest (XGRF), and MLP. This ensemble leverages the strengths of each model to enhance predictive performance and robustness. The overall performance on PRA201 is shown in Tab. 5. Notably, our goal is not to achieve SOTA performance on PRA201 but rather to design a robust affinity prediction model for our high-throughput screening framework.

**Table 5.** 5-fold regression performance of our affinity predictor on PRA201.

| Fold | MAE | RMSE | PCC | SCC |
| --- | --- | --- | --- | --- |
| 0 | 1.24 | 1.61 | 0.60 | 0.59 |
| 1 | 1.30 | 1.66 | 0.55 | 0.50 |
| 2 | 1.02 | 1.23 | 0.51 | 0.53 |
| 3 | 1.54 | 1.84 | 0.29 | 0.39 |
| 4 | 1.18 | 1.47 | 0.60 | 0.63 |
| Average | 1.26 | 1.56 | 0.51 | 0.53 |

## D. Evaluation Metrics

During evaluation, we leverage recovery rate and macro F1 as evaluation metrics following RDesign (Tan et al., 2024). As pointed out in RDesign, although perplexity is a crucial metric which has been widely used in protein design, its suitability is significantly less in RNA design. In RNA design scenario, the theoretical perplexity value for random guessing is 4, given the extremely small alphabet size {A, U, C, G}. However, in practical evaluations, most methods tend to exhibit a perplexity higher than 4, which might seem counterintuitive. This discrepancy occurs because perplexity is a metric designed for natural language processing (NLP) tasks with large vocabularies, whereas in RNA design, the extremely limited alphabet makes it less reflective of the true design objectives of perplexity. Thus, we evaluate our CARD and other comparison methods using recovery rate and macro F1.

Recovery rate is the nucleotide-level accuracy for a designed sequence *S* with length *L* compared to the ground truth sequence *T*.

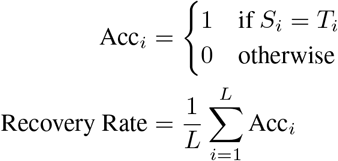

While macro F1 score provides a comprehensive measure of performance across different classes other than accuracy. This metric effectively captures the proportion of correctly classified instances at the letter level while balancing precision and recall, making it particularly useful when dealing with imbalanced classes.

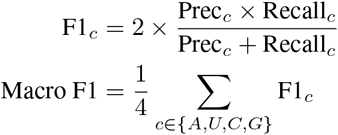

## E. Additional Case Study for High-Affinity RNA Design

In this section, we present more case studies using our CARD for high-affinity RNA sequence design.

### E.1. Design for 2HGH

2HGH is the complex of transcription factor IIIA (TFIIIA) zinc fingers 4-6 bound to 5S rRNA 55mer, while TFIIIA is a Cys2His2 zinc finger protein that regulates the expression of the 5 S ribosomal RNA gene. Similar to the process presented in Sec. 5, we input the 2HGH complex into CARD to perform RNA inverse folding to generate 1000 RNA sequences, and then predict their binding affinity. Next, we select the top 20 sequences with the highest predicted affinity and use AlphaFold3 to predict the structures, enabling a second-round design iteration. Through this iterative process, we generate 3 rounds of RNA sequences, progressively optimizing binding affinity for enhanced molecular interaction. We first show the sequence logo of generated sequences in Fig. 12. Although the RNA sequence length in 2HGH is longer compared to 2LBS, it retains more conserved regions. We conjecture that the structural constraints in 2HGH may lead to a higher degree of sequence preservation despite its increased length. The same phenomenon can also be shown in the affinity distribution of generated sequences across 3 rounds in Fig. 13. As observed in the figure, during the first two rounds, the high-affinity designs generated by CARD have already approached saturation. Consequently, in the third round, the newly generated sequences do not exhibit further improvements in affinity, and their distribution remains similar to that of the second round.

**Figure 12.**
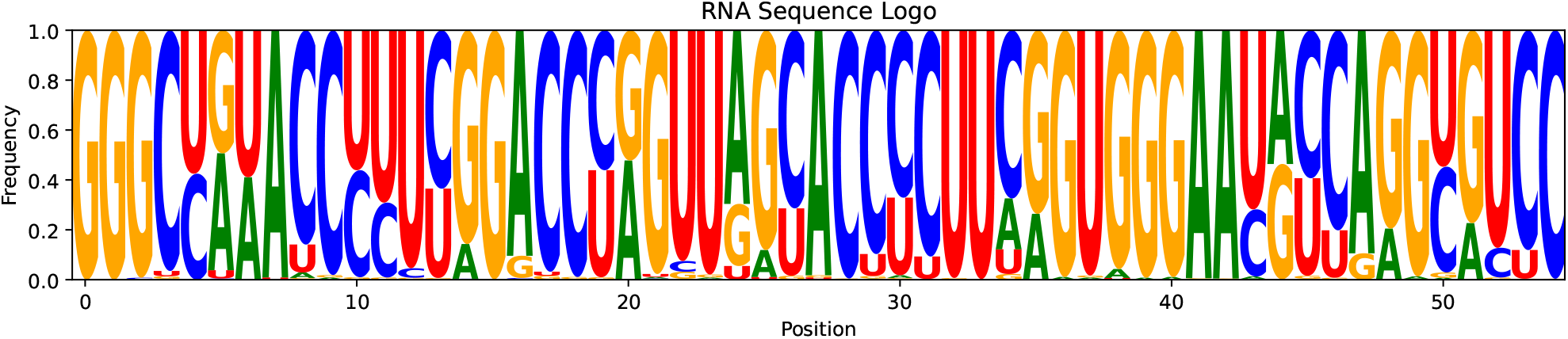
Sequence logo of generated RNA sequences for 2HGH

**Figure 13.**
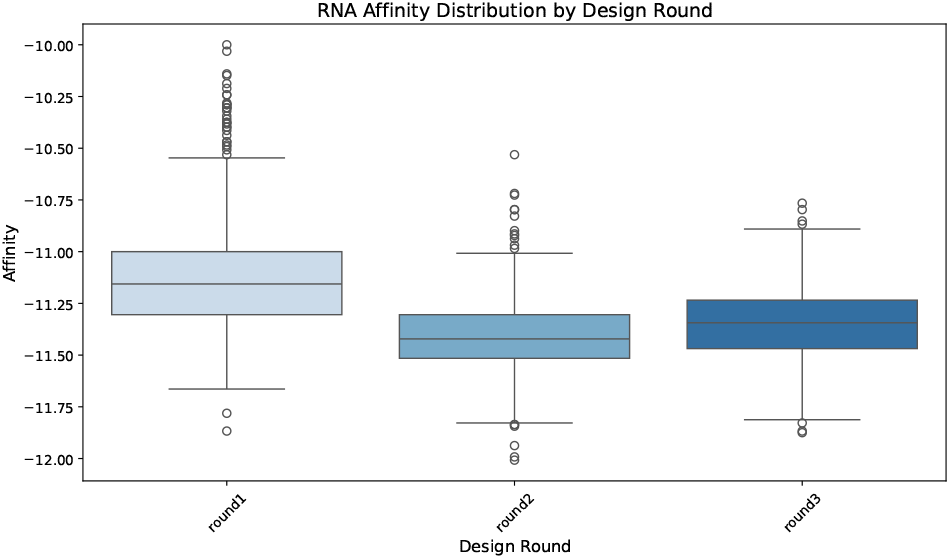
Affinity distribution of generated RNA sequences across round.

Here, we select two candidates from round 2 and round 3 with the best trade-off between structural constraint and affinity improvement. The structure is predicted by AlphaFold3. The predicted structures and generated sequences are shown in Fig. 14.

**Figure 14.**
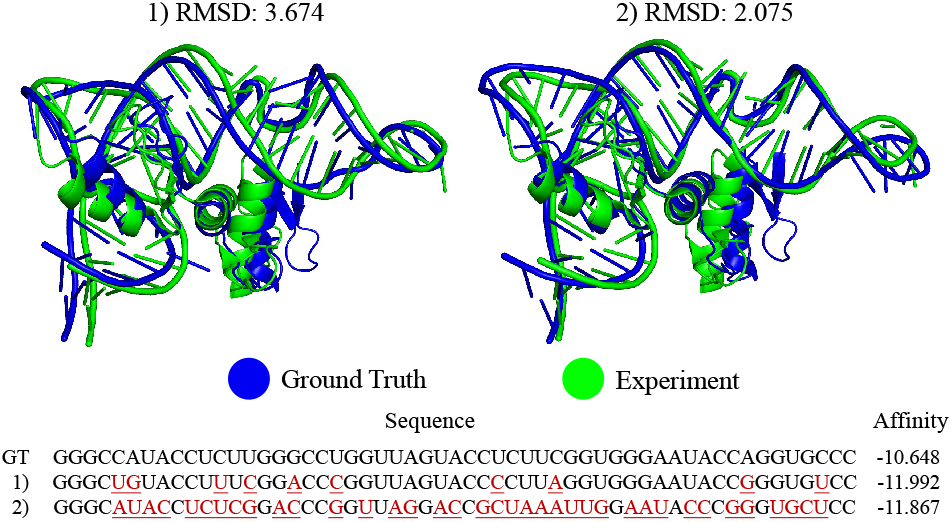
Structures and sequences of RNAs designed for 2HGH.

## References

Abramson, J., Adler, J., Dunger, J., Evans, R., Green, T., Pritzel, A., Ronneberger, O., Willmore, L., Ballard, A. J., Bambrick, J., et al. Accurate structure prediction of biomolecular interactions with alphafold 3. Nature, pp. 1–3, 2024.

Baek, M., McHugh, R., Anishchenko, I., Jiang, H., Baker, D., and DiMaio, F. Accurate prediction of protein–nucleic acid complexes using rosettafoldna. Nature methods, 21 (1):117–121, 2024.

Deng, L., Yang, W., and Liu, H. Predprba: prediction of protein-rna binding affinity using gradient boosted regression trees. Frontiers in genetics, 10:637, 2019.

Han, R., Liu, X., Pan, T., Xu, J., Wang, X., Lan, W., Li, Z., Wang, Z., Song, J., Wang, G., et al. Copra: Bridging cross-domain pretrained sequence models with complex structures for protein-rna binding affinity prediction. arXiv preprint arXiv:2409.03773, 2024.

Harini, K., Srivastava, A., Kulandaisamy, A., and Gromiha, M. M. Pronab: database for binding affinities of protein– nucleic acid complexes and their mutants. Nucleic acids research, 50(D1):D1528–D1534, 2022.

Harini, K., Sekijima, M., and Gromiha, M. M. Pra-pred: Structure-based prediction of protein-rna binding affinity. International Journal of Biological Macromolecules, 259: 129490, 2024.

Hayes, T., Rao, R., Akin, H., Sofroniew, N. J., Oktay, D., Lin, Z., Verkuil, R., Tran, V. Q., Deaton, J., Wiggert, M., et al. Simulating 500 million years of evolution with a language model. Science, pp. eads0018, 2025.

Hong, X., Tong, X., Xie, J., Liu, P., Liu, X., Song, Q., Liu, S., and Liu, S. An updated dataset and a structure-based prediction model for protein–rna binding affinity. Proteins: Structure, Function, and Bioinformatics, 91(9): 1245–1253, 2023.

Huang, H., Lin, Z., He, D., Hong, L., and Li, Y. Ribodif-fusion: tertiary structure-based rna inverse folding with generative diffusion models. Bioinformatics, 40:i347.–i356, 2024.

Ingraham, J., Garg, V., Barzilay, R., and Jaakkola, T. Generative models for graph-based protein design. Advances in neural information processing systems, 32, 2019.

Jing, B., Eismann, S., Suriana, P., Townshend, R. J., and Dror, R. Learning from protein structure with geometric vector perceptrons. arXiv preprint arXiv:2009.01411, 2020.

Joshi, C. K., Jamasb, A. R., Viñas, R., Harris, C., Mathis, S., Morehead, A., Anand, R., and Liò, P. grnade: Geometric deep learning for 3d rna inverse design. bioRxiv, 2024.

Keene, J. D., Komisarow, J. M., and Friedersdorf, M. B. Rip-chip: the isolation and identification of mrnas, micrornas and protein components of ribonucleoprotein complexes from cell extracts. Nature protocols, 1(1):302–307, 2006.

Ken, M. L., Roy, R., Geng, A., Ganser, L. R., Manghrani, A., Cullen, B. R., Schulze-Gahmen, U., Herschlag, D., and Al-Hashimi, H. M. Rna conformational propensities determine cellular activity. Nature, 617(7962):835–841, 2023.

Konig, J., Zarnack, K., Rot, G., Curk, T., Kayikci, M., Zupan, B., Turner, D. J., Luscombe, N. M., and Ule, J. iclip-transcriptome-wide mapping of protein-rna interactions with individual nucleotide resolution. Journal of visualized experiments: JoVE, (50):2638, 2011.

Lambert, N. J., Robertson, A. D., and Burge, C. B. Rna bind-n-seq: measuring the binding affinity landscape of rna-binding proteins. In Methods in enzymology, volume 558, pp. 465–493. Elsevier, 2015.

Li, W. and Godzik, A. Cd-hit: a fast program for clustering and comparing large sets of protein or nucleotide sequences. Bioinformatics, 22(13):1658–1659, 2006.

Licatalosi, D. D., Mele, A., Fak, J. J., Ule, J., Kayikci, M., Chi, S. W., Clark, T. A., Schweitzer, A. C., Blume, J. E., Wang, X., et al. Hits-clip yields genome-wide insights into brain alternative rna processing. Nature, 456(7221): 464–469, 2008.

Lin, Z., Akin, H., Rao, R., Hie, B., Zhu, Z., Lu, W., dos Santos Costa, A., Fazel-Zarandi, M., Sercu, T., Candido, S., et al. Language models of protein sequences at the scale of evolution enable accurate structure prediction. BioRxiv, 2022:500902, 2022.

Lorenz, R., Bernhart, S. H., Höner zu Siederdissen, C., Tafer, H., Flamm, C., Stadler, P. F., and Hofacker, I. L. Viennarna package 2.0. Algorithms for molecular biology, 6:1–14, 2011.

Pandey, U., Behara, S. M., Sharma, S., Patil, R. S., Nambiar, S., Koner, D., and Bhukya, H. Deepnap: A deep learning method to predict protein–nucleic acid binding affinity from their sequences. Journal of Chemical Information and Modeling, 64(6):1806–1815, 2024.

Runge, F., Stoll, D., Falkner, S., and Hutter, F. Learning to design rna. arXiv preprint arXiv:1812.11951, 2018.

Shen, T., Hu, Z., Sun, S., Liu, D., Wong, F., Wang, J., Chen, J., Wang, Y., Hong, L., Xiao, J., et al. Accurate rna 3d structure prediction using a language model-based deep learning approach. Nature Methods, pp. 1–12, 2024.

Tan, C., Zhang, Y., Gao, Z., Hu, B., Li, S., Liu, Z., and Li, S. Z. Rdesign: Hierarchical data-efficient representation learning for tertiary structure-based rna design. In The Twelfth International Conference on Learning Representations, 2024.

Tuerk, C. and Gold, L. Systematic evolution of ligands by exponential enrichment: Rna ligands to bacteriophage t4 dna polymerase. science, 249(4968):505–510, 1990.

Ule, J., Jensen, K., Mele, A., and Darnell, R. B. Clip: a method for identifying protein–rna interaction sites in living cells. Methods, 37(4):376–386, 2005.

Van Nostrand, E. L., Pratt, G. A., Shishkin, A. A., Gelboin-Burkhart, C., Fang, M. Y., Sundararaman, B., Blue, S. M., Nguyen, T. B., Surka, C., Elkins, K., et al. Robust transcriptome-wide discovery of rna-binding protein binding sites with enhanced clip (eclip). Nature methods, 13 (6):508–514, 2016.

Wang, R., Fang, X., Lu, Y., and Wang, S. The pdbbind database: Collection of binding affinities for protein-lig- and complexes with known three-dimensional structures. Journal of medicinal chemistry, 47(12):2977–2980, 2004.

Wong, F., He, D., Krishnan, A., Hong, L., Wang, A. Z., Wang, J., Hu, Z., Omori, S., Li, A., Rao, J., et al. Deep generative design of rna aptamers using structural predictions. Nature Computational Science, pp. 1–11, 2024.

Yang, W. and Deng, L. Pnab: prediction of protein-nucleic acid binding affinity using heterogeneous ensemble models. In 2019 IEEE International Conference on Bioinformatics and Biomedicine (BIBM), pp. 58–63. IEEE, 2019.

Yesselman, J. D. and Das, R. Rna-redesign: a web server for fixed-backbone 3d design of rna. Nucleic Acids Research, 43(W1):W498–W501, 2015.

Zhao, W. Ling-ling chen: Rna has its own features; don’t study it as a protein. National Science Review, 11(2): nwad287, 2024.

Zuker, M. Mfold web server for nucleic acid folding and hybridization prediction. Nucleic acids research, 31(13): 3406–3415, 2003.

